# Uncertainty quantified discovery of chemical reaction systems via Bayesian scientific machine learning

**DOI:** 10.1101/2023.09.11.557164

**Authors:** Emily Nieves, Raj Dandekar, Chris Rackauckas

## Abstract

The recently proposed Chemical Reaction Neural Network (CRNN) discovers chemical reaction pathways from time resolved species concentration data in a deterministic manner. Since the weights and biases of a CRNN are physically interpretable, the CRNN acts as a digital twin of a classical chemical reaction network. In this study, we employ a Bayesian inference analysis coupled with neural ordinary differential equations (ODEs) on this digital twin to discover chemical reaction pathways in a probabilistic manner. This allows for estimation of the uncertainty surrounding the learned reaction network. To achieve this, we propose an algorithm which combines neural ODEs with a preconditioned stochastic gradient langevin descent (pSGLD) Bayesian framework, and ultimately performs posterior sampling on the neural network weights. We demonstrate the successful implementation of this algorithm on several reaction systems by not only recovering the chemical reaction pathways but also estimating the uncertainty in our predictions. We compare the results of the pSGLD with that of the standard SGLD and show that this optimizer more efficiently and accurately estimates the posterior of the reaction network parameters. Additionally, we demonstrate how the embedding of scientific knowledge improves extrapolation accuracy by comparing results to purely data-driven machine learning methods. Together, this provides a new framework for robust, autonomous Bayesian inference on unknown or complex chemical and biological reaction systems.

## 2 Introduction

Mechanistic models in Quantitative Systems Pharmacology (QSP) offer more interpretability than purely data-driven machine learning approaches, but practitioners often lack complete knowledge of the underlying systems. Methods of Scientific Machine Learning (SciML) account for this epistemic uncertainty by mixing neural network techniques with mechanistic modeling forms to allow for the automated discovery of missing components in mechanistic models [1, 2]. In addition to interpretability, SciML models also offer superior prediction extrapolation beyond the domain in which they were trained, making them attractive candidates for QSP and systems biology applications.

The Chemical Reaction Neural Network (CRNN) is a recently proposed SciML method aimed at discovering chemical reaction pathways from concentration time course data without prior knowledge of the underlying system [3]. Specifically, the CRNN is a Neural ODE (Ordinary Differential Equation) architecture in that the neural network learns and represents the system of ODEs that comprise the reaction network [4]. The CRNN is a neural network architecture which is designed to be a flexible representation of mass action kinetics models, thus imposing prior known constraints about the guiding equations of chemical reaction models while allowing for interpretable weights which represent the individual reaction weights. The weights of the embedded neural networks represent the stoichiometry of the chemical species and kinetic parameters of the reactions. Optimization of the neural network weights thus leads to the discovery of the reaction pathways involved in the network. As QSP models often involve complex, interconnected reaction networks such as signalling or metabolic reactions, the CRNN may aid in the development of these models when the reactions are unknown or incompletely known.

The learned reaction system and the kinetic parameters of the CRNN are obtained in a deterministic manner, however to build trust in predictions and help experts understand when more training data is needed, the uncertainty of the predictions should also be quantified. For this purpose, Bayesian inference frameworks can be integrated with the CRNN. Bayesian methods obtain estimates of the posterior probability density function of neural network parameters allowing for an understanding of prediction uncertainty. It is the goal of this work to extend the CRNN with a Bayesian framework to increase its utility in QSP and other scientific domains. This is similar to the work by Li et al. in which they also extend the CRNN to include a Bayesian framework [5], but here we use a different methodology for sampling the posterior.

Recently, there has been an emergence of efficient Bayesian inference methods suitable for high-dimensional parameter systems, specifically methods like the No U-Turn Sampler (NUTS) [6], Stochastic Gradient Langevin Descent (SGLD) [7] and Stochastic Gradient Hamiltonian Monte Carlo (SGHMC) [8]. These methods are based on Markov Chain Monte Carlo (MCMC) sampling and utilize gradient information to obtain estimates of the posterior. A number of studies have explored the use of these Bayesian methods to infer parameters of systems defined by ODEs [9, 10, 11, 12], while others have used Bayesian methods to infer parameters of neural network models [13, 14, 15].

Specifically for Neural ODEs, the addition of Bayesian methods presents technical complexity as the training of the neural network is also tied to the solving of the ODE system. However recent work has demonstrated the feasibility of this approach [16]. To apply a Bayesian framework to a Neural ODE model, the authors make use of the Julia programming language which allows differential equation solvers [17, 18] to be combined with Julia’s probabilistic programming ecosystem [19, 20]. In this study, we similarly leverage the differential and probabilistic programming ecosystem of the Julia programming language to integrate the CRNN with Bayesian inference frameworks.

While powerful in their ability to estimate the posterior distribution, Bayesian inference methods may suffer from inefficient sampling of the posterior. The inefficiency often stems from the pathological curvature of the parameter space that leads to “bouncing” around minima. To address this, a new optimizer was proposed by Li et al. that combines an adaptive preconditioner with the SGLD algorithm [21]. The preconditioner performs a local transform of the parameter space such that the rate of curvature is the same in all directions, allowing for smoother, faster descent.

In this study, we propose combining the CRNN with the more efficient preconditioned SGLD optimizer [21] to discover chemical reaction networks and quantify the uncertainty of the learned reaction parameters, allowing for its more robust use in QSP and other scientific domains. We apply this Bayesian SciML method to a systems biology pathway to demonstrate its application to QSP. We also compare the results to those generated using purely data-driven machine learning methods to further demonstrate the advantages of SciML methods.

## 3 Methodology

Similarly to the originally proposed CRNN [3], we define the Chemical Reaction Neural Network (CRNN) to represent the following elementary reaction:

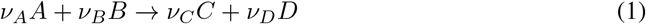

where *ν* refers to the stoichometric coefficients of the respective chemical species.

The law of mass action applied to the example reaction prescribed in equation 1 leads to the following expression for the reaction rate *r*:

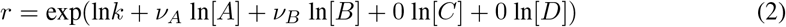

where k is the rate constant of the reaction and [A] refers to the concentration of chemical species A.

This elementary reaction can be represented by a neuron governed by *y* = *σ*(*wx* + *b*) where *w* are the weights, *y* is the neuron output, *x* is the input to the neuron and *σ* is the activation function. The inputs are the concentration of the species in logarithmic scale, and the output is the production rate of all species: 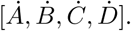 The input layer weights denote the reaction orders i.e [*ν*_*A*_, *ν*_*B*_, 0, 0] for the reactants [*A, B, C, D*] respectively. The output layer weights denote the stochiometric coefficients i.e [−*ν*_*A*_, *−ν*_*B*_, *ν*_*C*_, *ν*_*D*_] for [*A, B, C, D*] respectively. The bias denotes the rate constant in the logarithmic scale. This is depicted in Figure 1.

**Figure 1.**
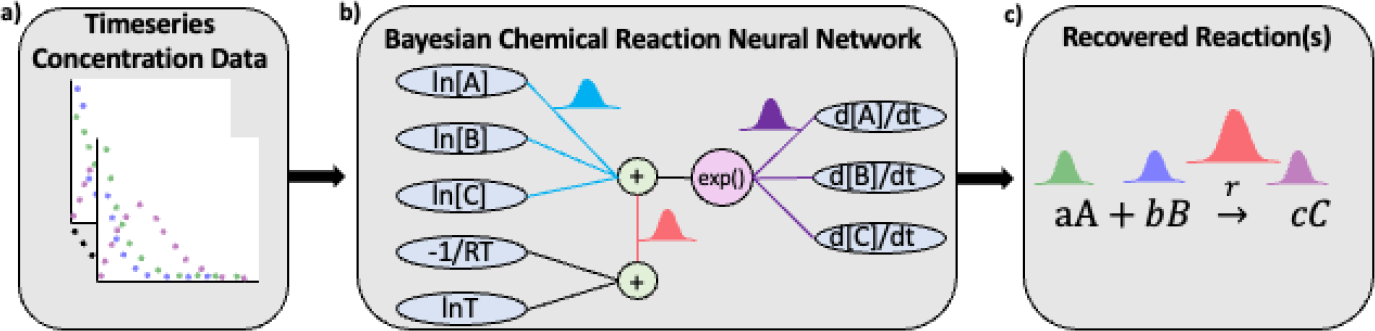
Overview of the Bayesian chemical reaction network which uses timeseries concentration data (a) to train a constrained neural network (b) that uses a preconditioned SGLD optimizer to reconstruct the reaction network and estimate the uncertainty in the learned stoichiometry and reaction rates (c).

A reaction network with multiple chemical reactions can be denoted by a CRNN with a hidden layer. Since the weights and biases of a CRNN are physically interpretable, it is a digital twin of a classical chemical reaction neural network. In this study, we employ Bayesian inference analysis on this digital twin to discover chemical reaction pathways in a probabilistic manner.

Considering the vector of chemical species *Y* to be varying in time, we aim to recover a CRNN which satisfies the following equation:

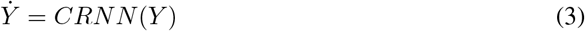

where 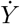 and is the rate of change of the concentration of chemical species Y.

Thus, by representing the ODE governing the variation of the chemical species *Y* by a neural network, we can solve equation 3 using powerful, purpose built ODE solvers provided by the differential programming ecosystem of the Julia programming language [18]. The solution we obtain by solving the ODE is denoted by *Y*_CRNN_(*t*). In this study, the loss function between the actual species concentrations and the predicted values is governed by the Mean Absolute Error (MAE) with L2 regularization, and given by

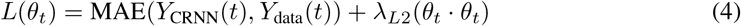

where *θ*_*t*_ are the weights of the CRNN at time *t*, λ_*L*2_ is the L2 regularization coefficient,*Y*_*CRNN*_ (*t*) are the CRNN predicted values of the species concentrations at time t, and *Y*_*data*_(*t*) is the true species concentration at time t.

To incorporate uncertainty into our predictions, we augment the gradient descent algorithms with a Bayesian framework. Our algorithm is shown in Algorithm 1. We use a variation of the Stochastic Gradient Langevin Descent (SGLD) algorithm [7] and add a preconditioning term *G*(*θ*_*t*_), which adapts to the local geometry leading to more efficient training of deep neural networks as noted in [21].

Similarly to [21], we utilize a decreasing step-size that’s given by the following equation:

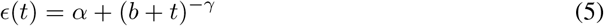

where ϵ(*t*) is the step-size, t is the epoch number, and *α*, b and *γ* are tunable parameters.

### Algorithm 1 Preconditioned SGLD applied to Neural ODE

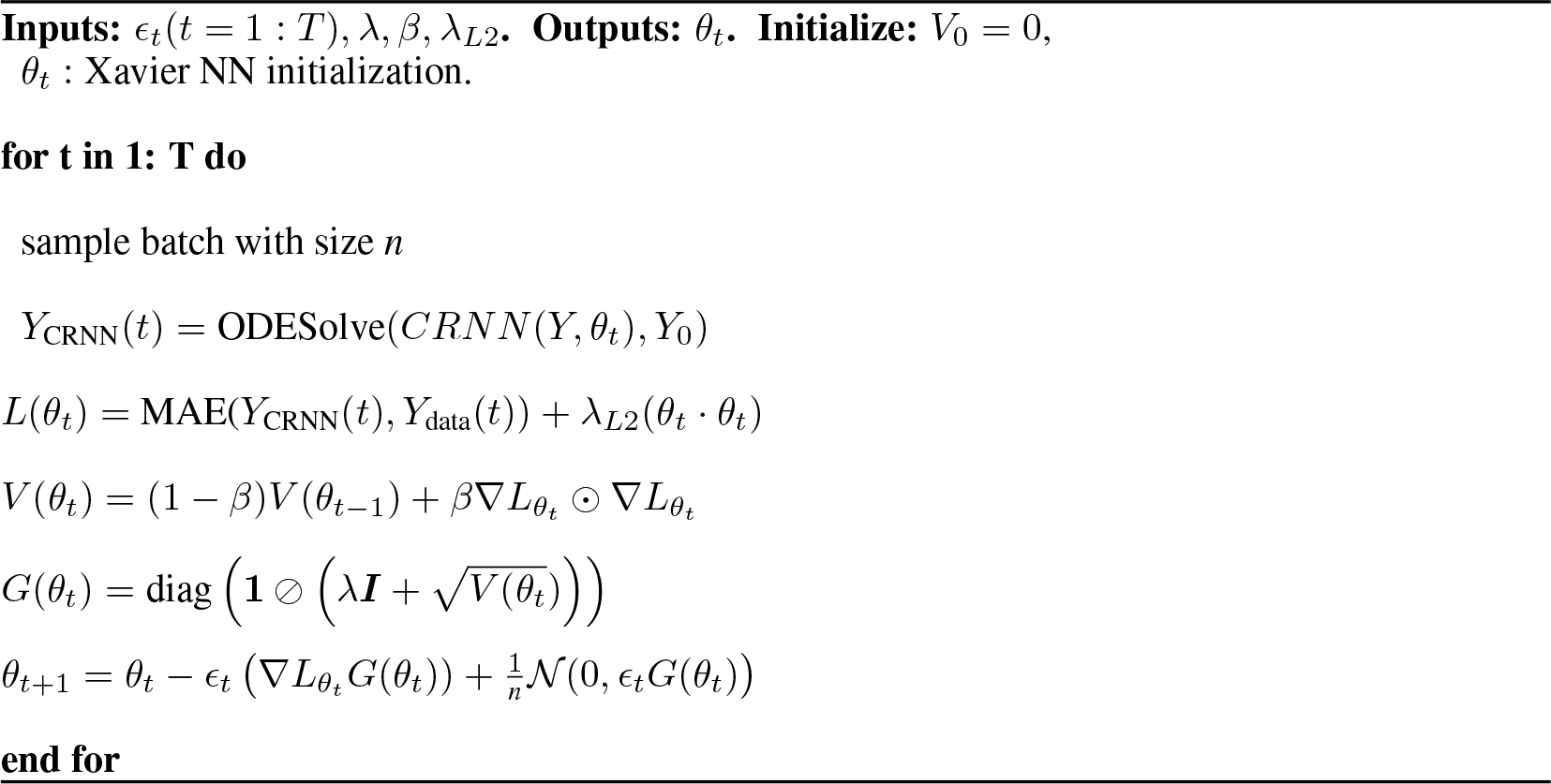

In this algorithm, the induced noise and step-size decay to zero as the training proceeds. By adding Gaussian noise to each step of the gradient descent, the optimizer finds a local model, but never converges due to the induced noise. The decayed learning rate also prevents the optimizer from leaving the model, leading to random walking around the mode. The process after settling in a local model is the sampling phase, and in this phase we draw parameter posterior samples. The point at which the optimizer enters the sampling phase can be determined by observing when the loss function stagnates. Thus, the parameters *θ* of the CRNN obtained during the sampling phase lead to the parameter posterior which ultimately helps in encoding uncertainty in the prediction of the chemical reaction pathways.

## 4 Results

### 4.1 Case 1: Simple reaction network

We first consider the chemical reaction given in [22] comprising of five species: [*A, B, C, D, E*] involved in four reactions, described in Table 1. Similar to [3], a total of 100 synthetic datasets were simulated with initial conditions randomly sampled between [0.2, 0.2, 0, 0, 0] and [1.2, 1.2, 0, 0, 0]. Each dataset is comprised of 100 time points. Out of the 100 datasets, 90 are used for training and the remaining are reserved for validation. Further, mini-batching is employed to accelerate the training process and add additional regularization. As the validation loss function stagnates, early stopping is employed. Mini-batching and early stopping prevent the CRNN from overfitting the training data. Here, we choose *λ* = 1*e* −6, *β* = 0.9, and *λ*_*L*2_ = 1*e* −5. For the step-size parameters we choose *α* = 0.001, *b* = 0.15, and *γ* = 0.005.

**Table 1:** Case 1 ground truth reactions Equation Rate.

| Equation | Rate |
| --- | --- |
| $2A \rightarrow B$ | 0.1 |
| $A \rightarrow C$ | 0.2 |
| $C \rightarrow D$ | 0.13 |
| $B + D \rightarrow E$ | 0.3 |

Figure 2 shows the comparison of the Bayesian Neural ODE prediction compared to the data for the species A, B, C, D, and E in the four reaction system described in Case 1. A total of 500 posterior sample prediction are superimposed on the data; and a good agreement is seen between all trajectories and the data.

**Figure 2.**
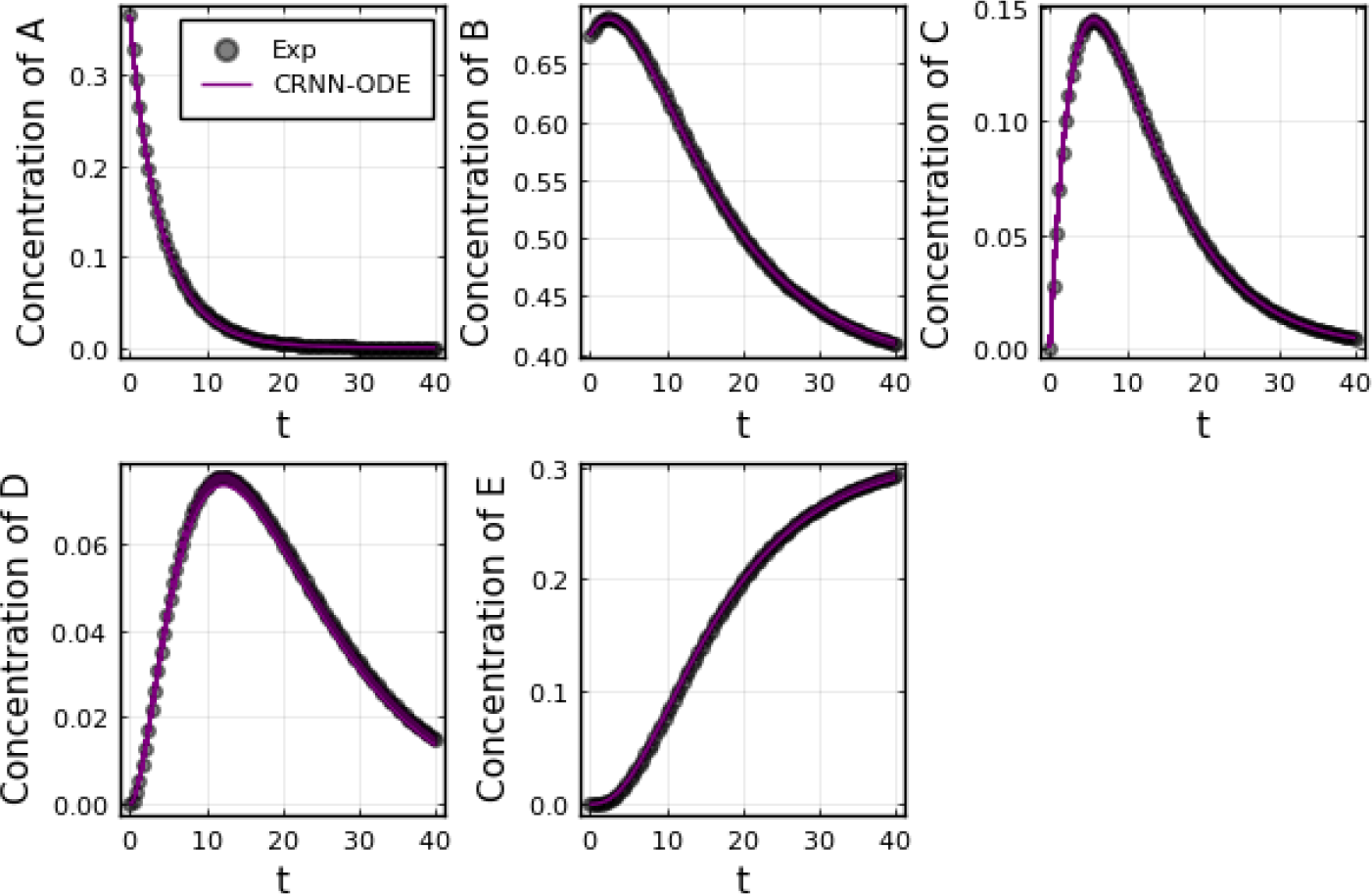
Comparison of the Bayesian chemical reaction neural network prediction to the data for the species A, B, C, D, and E in the four reactions described in Case 1. No noise was added to the data. A total of 500 posterior sample predictions are superimposed on the data.

Figure 3 shows the recovery probability of species A, B, C, D, and E in the four reactions described in Case 1, obtained using the preconditioned SGLD described in Algorithm 1. A posterior set of 1000 samples was chosen for the estimation. A species is considered to be present in a particular reaction if its weight is greater than 1*e* −4. The probability a species is contributing to a reaction is given by the ratio of the number of samples in which the species is present in that reaction to the total number of posterior samples (1000 in this case).

**Figure 3.**
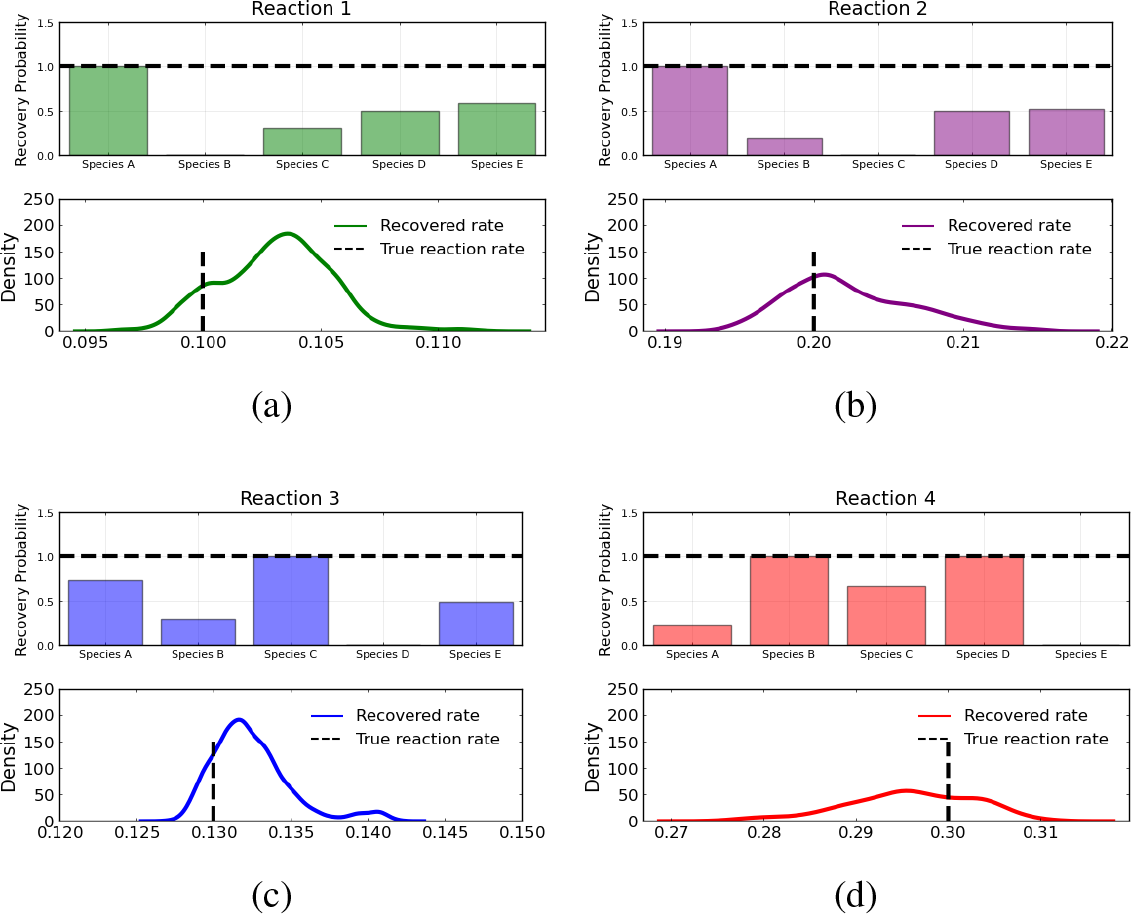
Reactant recovery probability of species A, B, C, D, and E and posterior distributions of learned reaction rates for the four reactions described in Case 1. No noise was added to the training data and a posterior set of 1000 samples was chosen for the estimation.

However, even if the probability of a species to be present in a reaction is high, its weight can be low compared to other species. Thus, we define a score metric for a species *i* in reaction *j* governed by weight *w*_*ij*_ to be as follows:

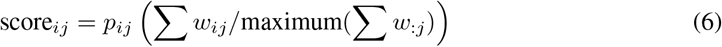

where *p*_*ij*_ is the recovery probability for the species shown in Figure 3, Σ *w*_*ij*_ is the summation of the weight values of species *i* for reaction *j* over all posterior samples and maximum(*w*_:*j*_) is the maximum value of this summation for the reaction *j* among all species.

From the recovered score metrics shown in Figure 4, we can see that we obtain a sparser discovery of the reaction pathways; which match with the data shown in Table 1.

**Figure 4.**
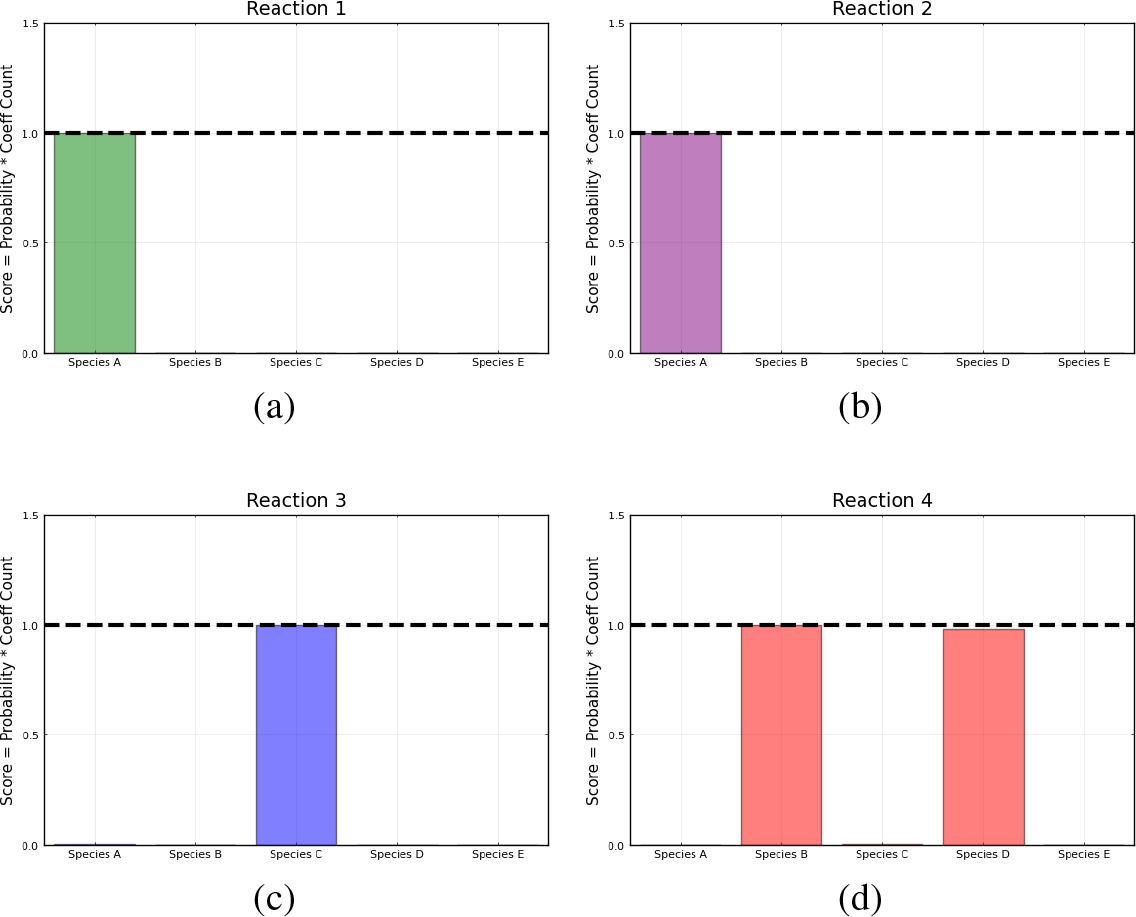
Recovered score metric for each of the four reactions shown in case 1. Score metrics are calculated using equation 6 and represent the weighted probability that a species is a reactant of each reaction. No noise was added to the data.

#### 4.1.1 Moderately noisy data

We subsequently train on moderately noisy data with standard deviation of the noise being 5% of the concentrations. The data is visualized in Figure 5. Figure 5 shows the comparison of the Bayesian Neural ODE predictions to the data for 500 posterior samples. Even with the addition of noise, our methodology can correctly capture the time course of all the chemical species.

**Figure 5.**
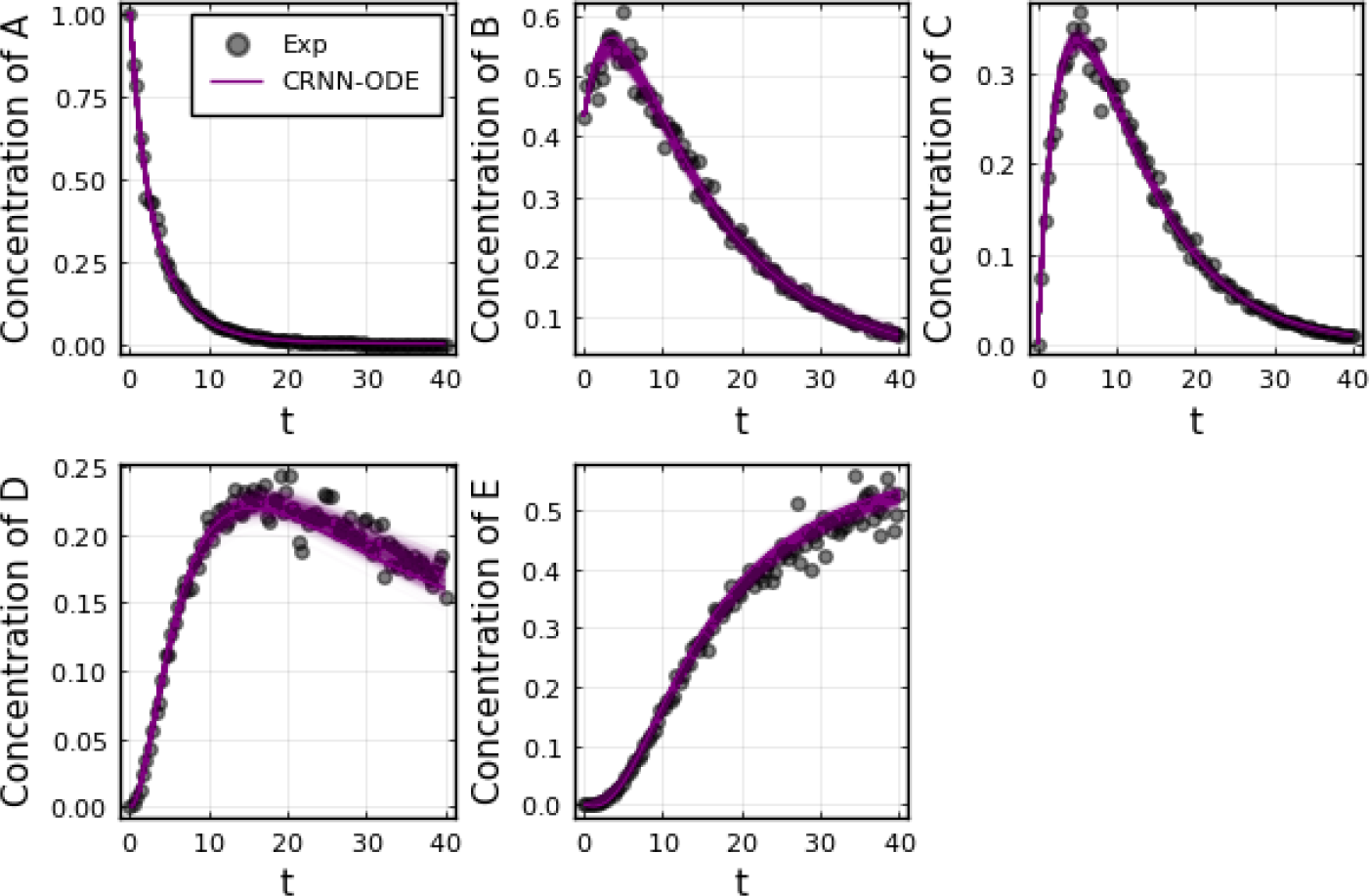
Comparison of the Bayesian chemical reaction neural network prediction to the data for the species A, B, C, D, and E in the four reactions described in Case 1. The model was trained with moderately noisy data, where the standard deviation of the noise was set to 5% of the concentrations. A total of 500 posterior sample predictions are superimposed on the data.

The reactant recovery probability and the score metric plots are shown in Figures 6, 7 respectively. Even with the presence of noise, the probability plots are seen to assign higher probability to the correct reaction pathways. The score plots accurately predict the reaction pathways as seen in the data (Table 1). Figure 6 also shows the posterior distribution of the reaction rates. Even with the addition of the noise, the distributions are centered close to the true reaction rate for all reactions except reaction 1, which still contains the true reaction rate within its distribution.

**Figure 6.**
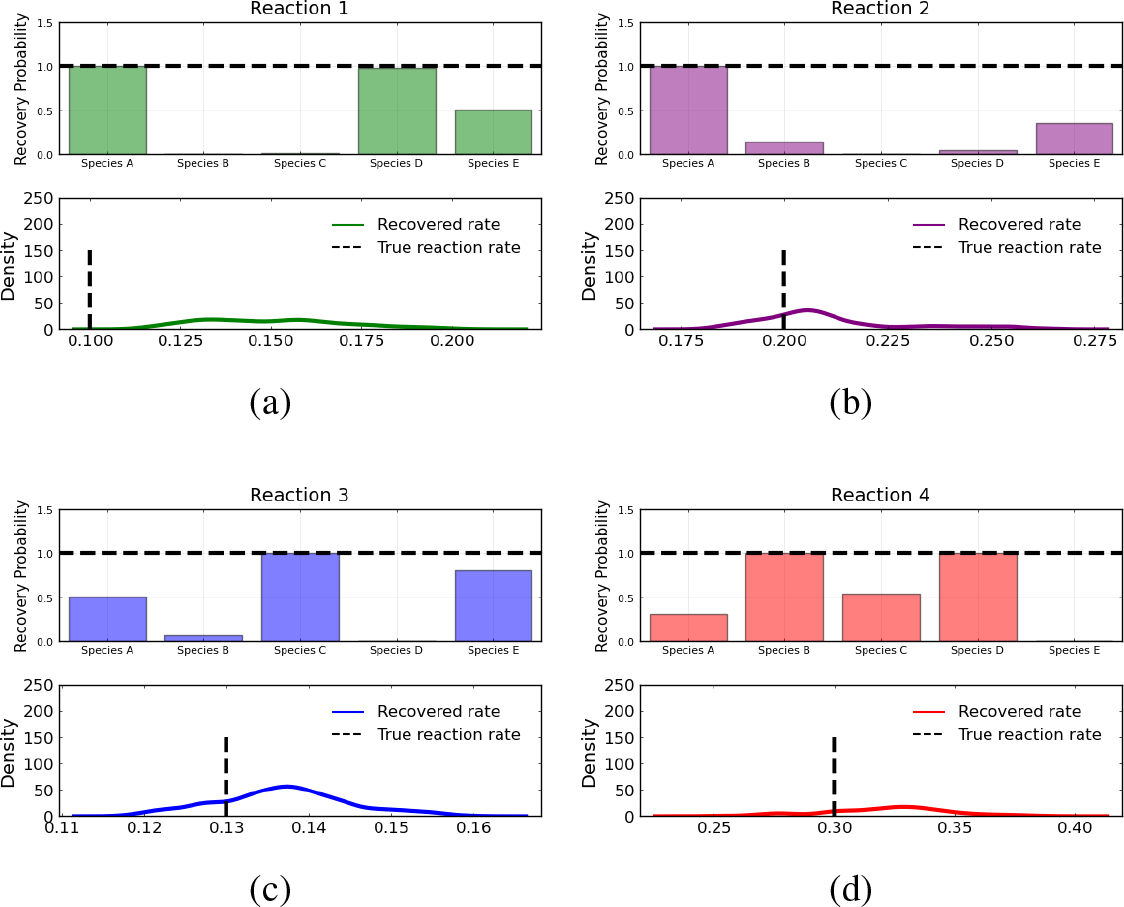
Reactant recovery probability of species A, B, C, D, and E and posterior distributions of learned reaction rates for the four reactions described in Case 1. The model was trained with moderately noisy data, where the standard deviation of the noise was set to 5% of the concentrations. A posterior set of 1000 samples was chosen for the estimation.

**Figure 7.**
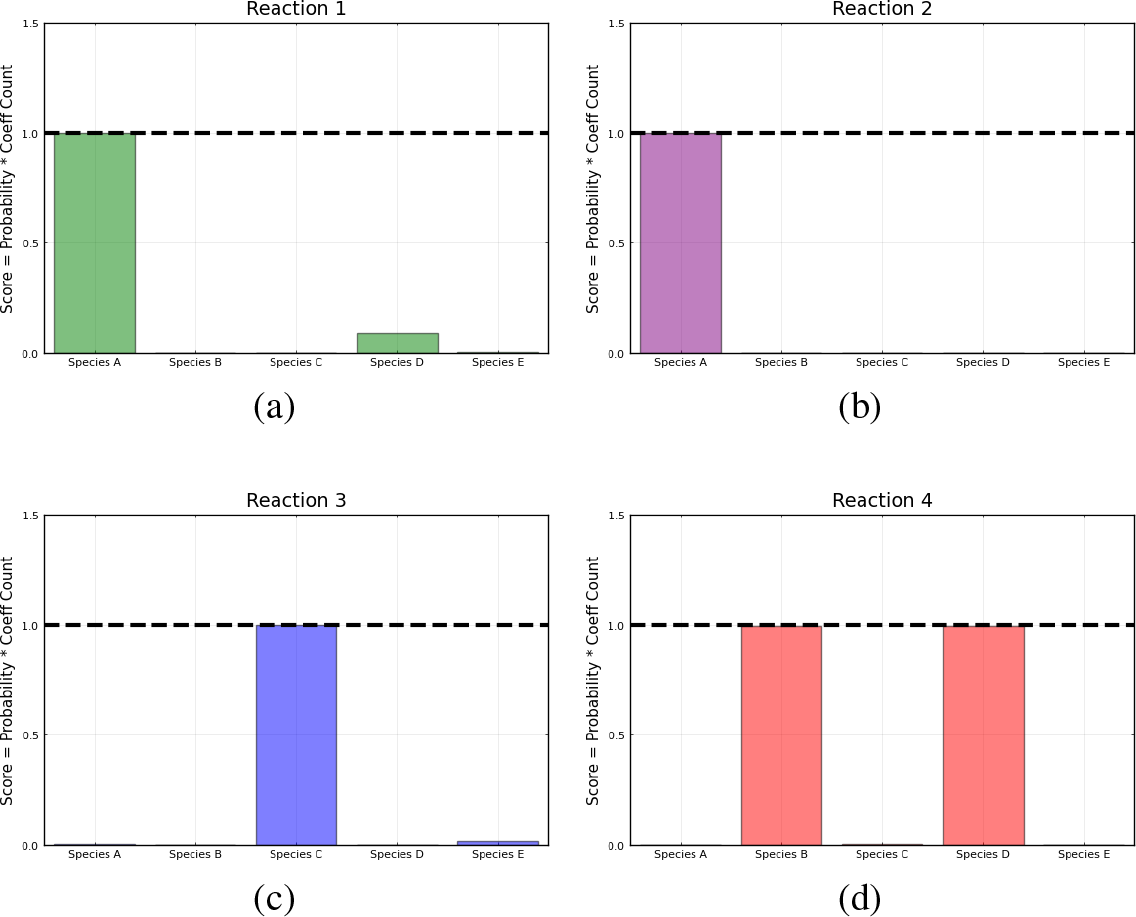
Recovered score metrics for each of the four reactions shown in case 1. Score metrics are calculated using equation 6 and represent the weighted probability that a species is a reactant of each reaction. The model was trained with moderately noisy data, where the standard deviation of the noise is set to 5% of the concentrations.

#### 4.1.2 Highly noisy data

To test the limits of the Bayesian Neural ODE framework, we subsequently train on highly noisy data with the standard deviation of the added noise being 50% of the concentrations. The Bayesian Neural ODE predictions are visualized in Figure 8 alongside the data. We observe that even with 50% noise, our methodology captures the data well. The added noise does however prevent the model from perfectly matching the true concentration time courses of each species.

**Figure 8.**
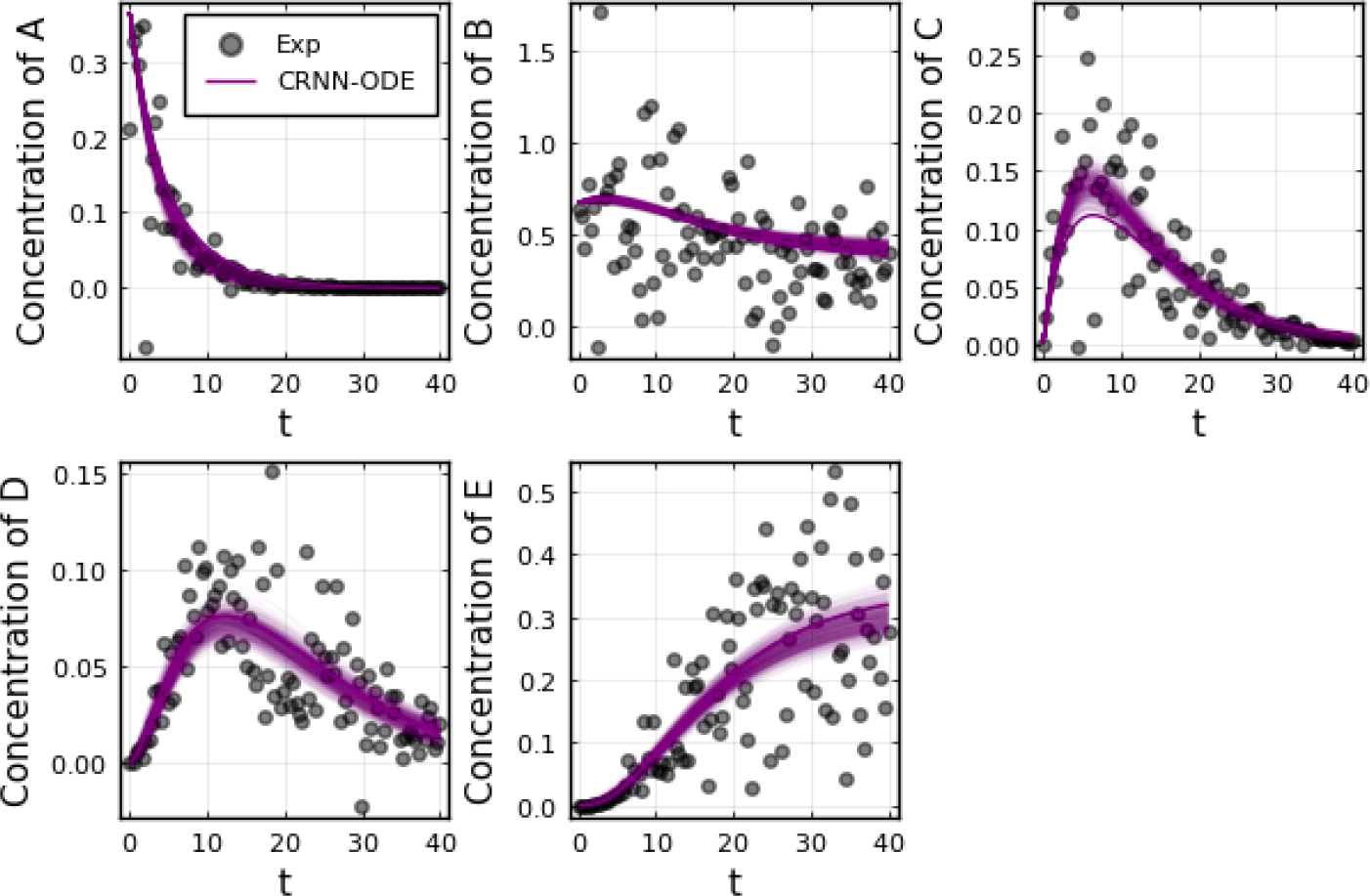
Comparison of the Bayesian chemical reaction neural network prediction compared to the data for the species A, B, C, D, and E in the four reactions described in Case 1. The model was trained with highly noisy data. The standard deviation of the noise is set to 50% of the concentrations. A total of 500 posterior sample predictions are superimposed on the data.

The reactant recovery probability and the score metric plots are shown in Figures 9, 10 respectively. Although a number of reactants show up in the probability plots due to the high amount of noise in the data, the score plots still accurately predict the reaction pathways as seen in the data (Table 1). Similarly, the posterior distributions of the reaction rates shown in Figure 9 contain the true reaction rates, however they are no longer centered at the true reaction rate, except for reaction 4.

**Figure 9.**
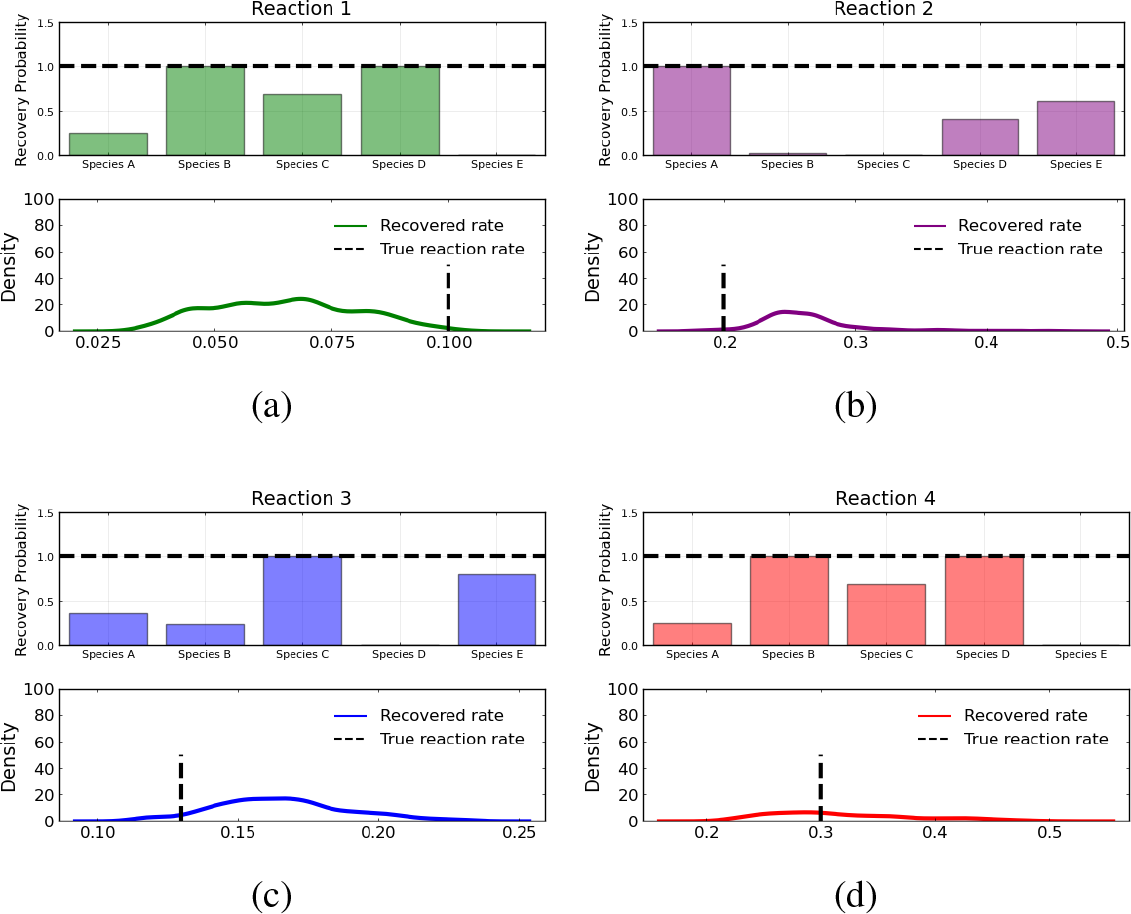
Reactant recovery probability of species A, B, C, D, and E and posterior distributions of learned reaction rates for the four reactions described in Case 1. The model was trained with highly noisy data. The standard deviation of the noise is set to 50% of the concentrations. A posterior set of 1000 samples was chosen for the estimation.

**Figure 10.**
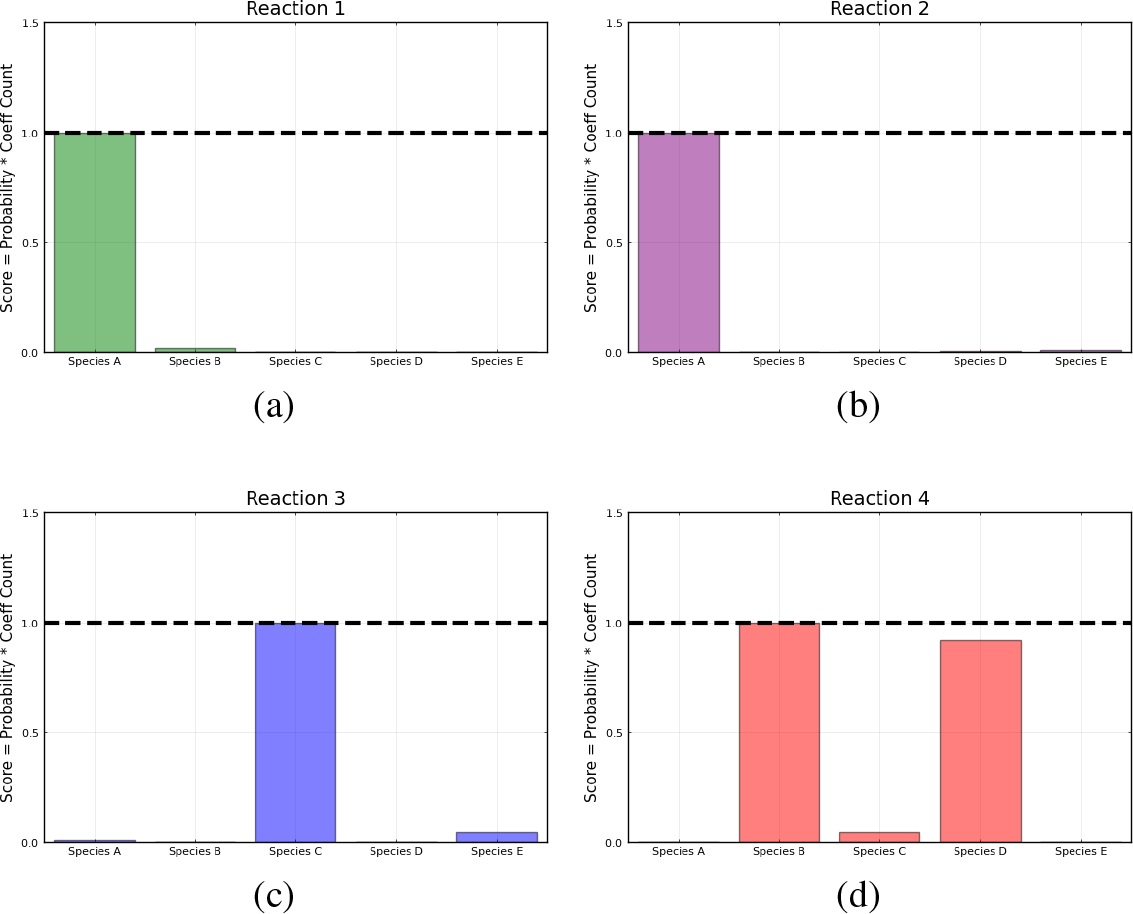
Recovered score metric for each of the four reactions shown in case 1. Score metrics are calculated using equation 6 and represent the weighted probability that a species is a reactant of each reaction. The model is trained with highly noisy data, where the standard deviation of the noise is set to 50% of the concentrations.

#### 4.1.3 Comparison to SGLD

We compare our results to the standard SGLD optimizer to demonstrate the effect of the preconditioner. Figure 11 shows training and validation loss over epochs for both the pSGLD and the standard SGLD. The preconditioned algorithm allows for faster entry into the sampling phase, entering around 6, 500 epochs before the SGLD.

**Figure 11.**
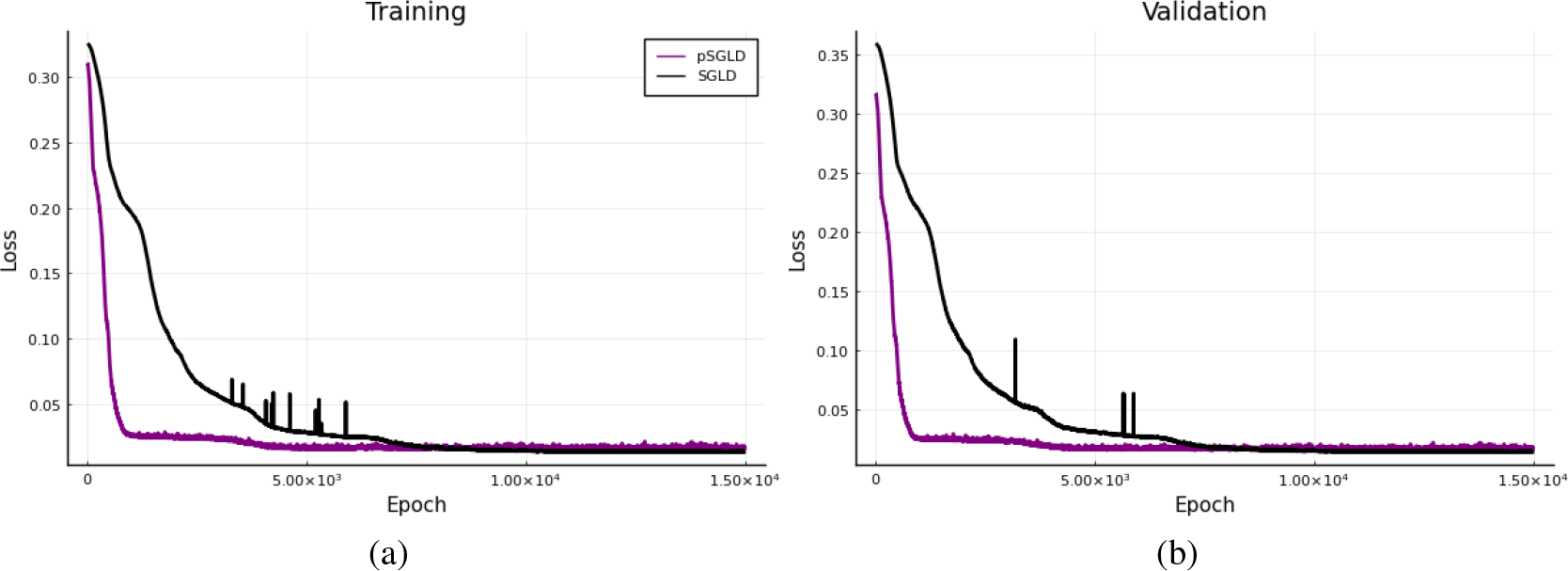
Comparison of training (a) and validation (b) losses over epochs with the preconditioned SGLD optimizer and SGLD optimizer for case 1 with 5% noise added.

Figure 12 compares the learned reaction rates in the case where 5% noise is added to the training data between the pSGLD and the SGLD. From this figure we can see that in general the pSGLD achieved a posterior that more closely matched the true reaction rates. The average percent deviation of the samples from the true reaction rates for the pSGLD were 50.49% ± 20.01, 7.44% ± 7.73, 6.70% ± 4.68, and 9.50% ± 6.14 for reactions 1, 2, 3, and 4 respectively. The average percent deviation from the true reaction rates for the standard SGLD were 3.78% ± 2.60, 6.96% ± 2.17, 13.27% ± 2.49, and 25.65% ± 4.52 for reactions 1, 2, 3, and 4 respectively. The pSGLD is similar or better in all reactions except for reaction 1. In reaction 1, the pSGLD has lower confidence, as shown in the low density, wide coverage of the posterior samples. In the other reactions, the SGLD demonstrates higher confidence in incorrect reaction rates whereas the pSGLD better represents the uncertainty by estimating a posterior that is centered at the true reaction rate.

**Figure 12.**
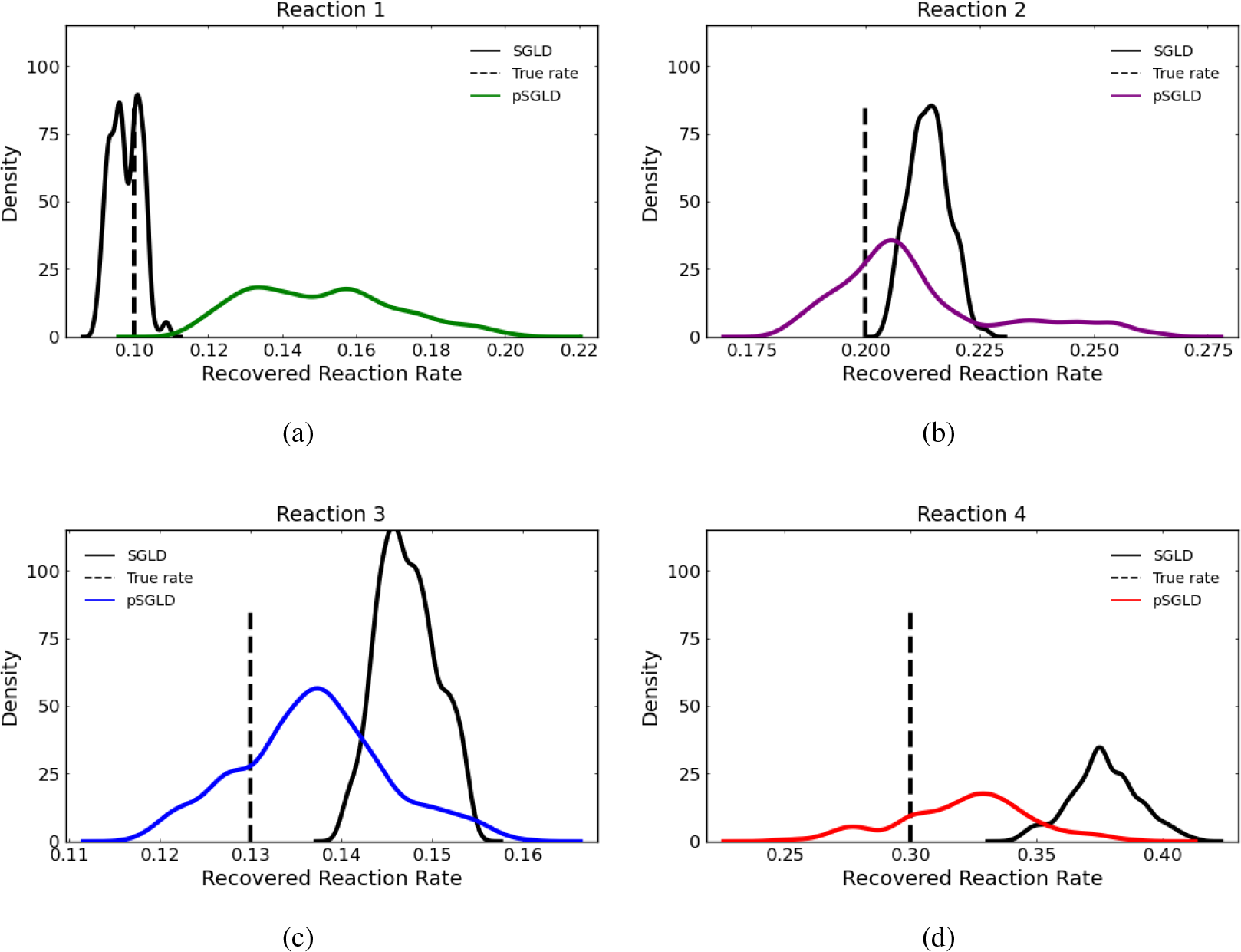
Comparison of estimated posterior of recovered reaction rates between the preconditioned SGLD and SGLD optimizer for the four reactions in case 1 where noise with a standard deviation of 5% was added.

**Figure 13.**
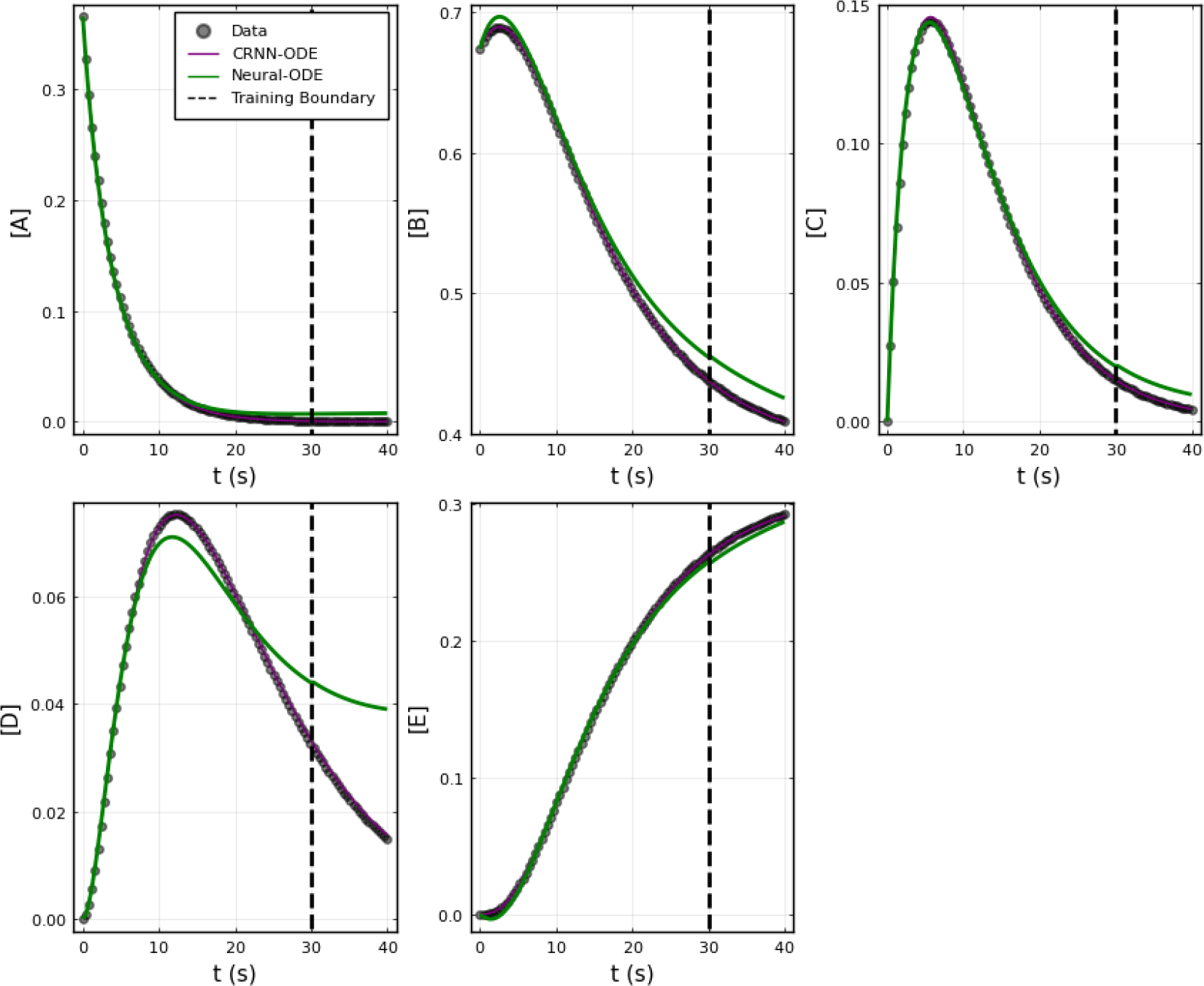
Comparison of Neural ODE with Bayesian CRNN (chemical reaction neural network) Only time points 0-30 were used for training as depicted by the training boundary. Time points 30-40 were used for comparison of the accuracy of extrapolation beyond training time points.

#### 4.1.4 Comparison to Neural ODE

We also compared the Bayesian CRNN to a purely data-driven neural ODE approach, where the structure does not include embedded scientific knowledge. For the neural ODE, we built a densely connected neural network with two hidden layers with 50 nodes each and hyperbolic tangent activation functions. The neural network was trained for 2000 epochs with the ADAM optimizer (learning rate set to 0.001). Here we compare the ability of the learned networks to extrapolate beyond the time points used for training. We use timepoints 0-30s for training and then use the trained system to predict concentrations 30-40 extrapolate to timepoint 40.

The results are summarized in Figure 13 and demonstrate the superior ability of the CRNN to accurately predict beyond region used for training. The mean absolute percent error in the extrapolation region was 9.0% ± 3.2 for the CRNN in comparison to 254.7% ± 109.8 for the neural ODE.

#### 4.1.5 Comparison to LSTM

Additionally, we compared our Bayesian CRNN to a more standard machine learning solution, in this case a Long Short-Term Memory (LSTM) model. This architecture predicts a sequence, in this case the timecourses for the chemical reaction species. We utilize the LSTM modules from Flux.jl [23], and connect two LSTM modules with 200 hidden nodes each to a densely connected linear output layer. Similarly to the neural ODE, we train for 2000 epochs with the ADAM optimizer (learning rate set to 0.001).

From Figure 14, we can see that the results of the CRNN and LSTM model are nearly indistinguishable in the training region, but the LSTM model diverges from the true concentration values in the extrapolation region while the CRNN matches the true values. The mean absolute percent error in the extrapolation region was 9.0% ± 3.2 for the CRNN in comparison to 90.93% ± 58.83 for the LSTM-based model.

**Figure 14.**
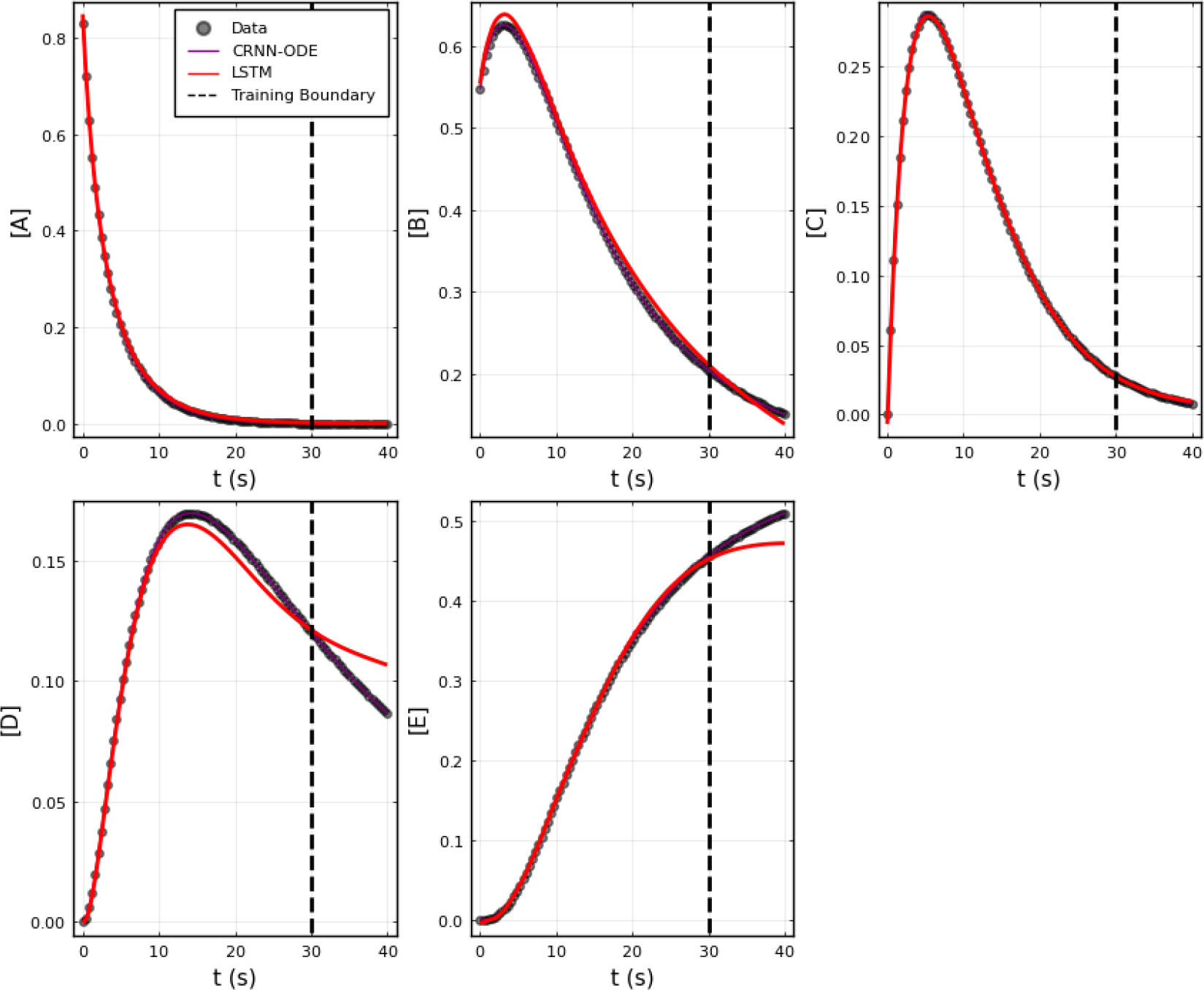
Comparison of LSTM model with Bayesian CRNN (chemical reaction neural network) Only time points 0-30 were used for training as depicted by the training boundary. Time points 30-40 were used for comparison of the accuracy of extrapolation beyond training time points.

### 4.2 Case 2: EGFR-STAT3 pathway

To further test the capabilities of the Bayesian CRNN, we attempted to recover the reaction network governing the EGFR-STAT3 signaling pathway [24]. Our simplified representation of this pathway consists of seven chemical species and six reactions listed in Table 2 and an overview is depicted in Figure 15. Here we omit the reverse reactions because they cannot be identified from the forward reactions with this framework.

**Table 2:**
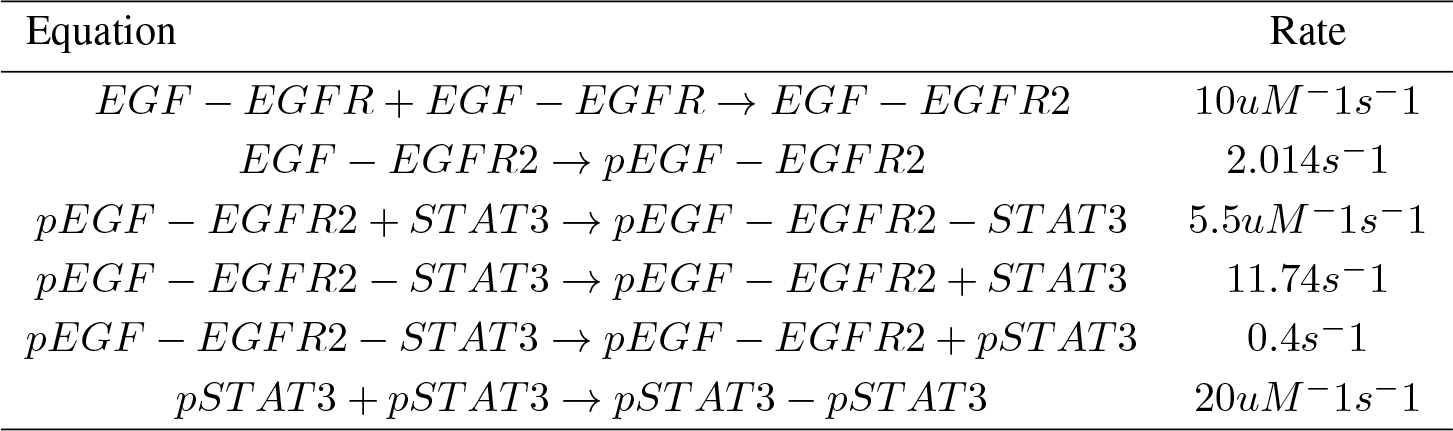
Case 2 EGFR-STAT3 reactions.

| Equation | Rate |
| --- | --- |
| $EGF - EGFR + EGF - EGFR \rightarrow EGF - EGFR2$ | $10uM^{-1}s^{-1}$ |
| $EGF - EGFR2 \rightarrow pEGF - EGFR2$ | $2.014s^{-1}$ |
| $pEGF - EGFR2 + STAT3 \rightarrow pEGF - EGFR2 - STAT3$ | $5.5uM^{-1}s^{-1}$ |
| $pEGF - EGFR2 - STAT3 \rightarrow pEGF - EGFR2 + STAT3$ | $11.74s^{-1}$ |
| $pEGF - EGFR2 - STAT3 \rightarrow pEGF - EGFR2 + pSTAT3$ | $0.4s^{-1}$ |
| $pSTAT3 + pSTAT3 \rightarrow pSTAT3 - pSTAT3$ | $20uM^{-1}s^{-1}$ |

**Table 3:** Average percent deviations from true reaction rates for case 2.

| Equation | Average Percent Deviation |
| --- | --- |
| $EGF - EGFR + EGF - EGFR \rightarrow EGF - EGFR2$ | $1.22\% \pm 0.81$ |
| $EGF - EGFR2 \rightarrow pEGF - EGFR2$ | $9.74\% \pm 0.64$ |
| $pEGF - EGFR2 + STAT3 \rightarrow pEGF - EGFR2 - STAT3$ | $22.41\% \pm 0.77$ |
| $pEGF - EGFR2 - STAT3 \rightarrow pEGF - EGFR2 + STAT3$ | $38.59\% \pm 0.56$ |
| $pEGF - EGFR2 - STAT3 \rightarrow pEGF - EGFR2 + pSTAT3$ | $643.32\% \pm 8.92$ |
| $pSTAT3 + pSTAT3 \rightarrow pSTAT3 - pSTAT3$ | $4.08\% \pm 1.28$ |

**Figure 15.**
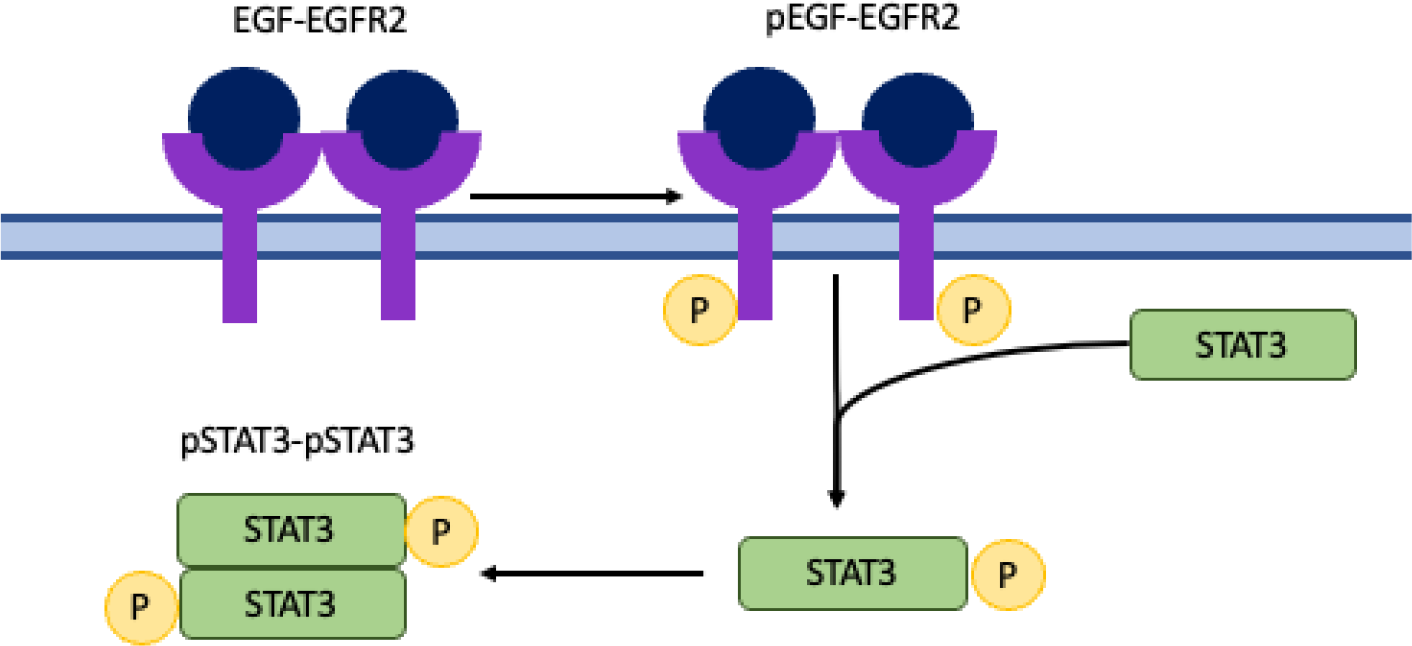
Simplified overview of EGFR-STAT3 reactions included in case 2, involving EGF-EGFR dimerization, EGF-EGFR dimer activation, STAT3 activation, and STAT3 dimerization.

In this case, unlike the simple system in case one, the reaction rates and concentrations vary by one or two orders of magnitude. To account for this difference in scale, we adjust our loss function to use mean absolute percent error (MAPE) instead of MAE. Additionally to allow for more reactions and chemical species, we use L1 regularization on the weights that correspond to the stoichiometric coefficients of the reactions *θ*_*ν*_, as these would comprise a sparse matrix as the reaction network grows. We continue to use L2 regularization for the weights that correspond to the reaction rates, *θ*_*r*_. This change to the loss function is depicted in the equation below.

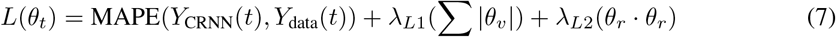

We sample from the initial conditions ranging from 0 to 1 to obtain 300 training examples. With 5% Gaussian noise added to simulated training data. For training, we set *λ* = 1*e* −8, *β* = 0.9, the L1 coefficient (λ_*L*1_) = 1e-4 and the L2 coefficient (λ_*L*2_) =1e-5. Step-size hyperparameters *α*, b, and *γ* were set to 0.001, 0.15, and 0.005 respectively.

After training, the Bayesian CRNN accurately captures the time course for all species in the simplified EGFR-STAT3 pathway and is robust to noise as shown in Figure 16 and from the score plots in Figure 17, we can see the CRNN also correctly identifies the reactants of all the reactions. Figure 18 shows the uncorrected reactant probabilities and the probability densities for the reaction rates. The true reaction rates are contained within the probability density functions of reactions 1 and 6 depicted in Figure 18 a and f, but the other reactions do not include the true rates despite the time courses of the reactants and products being accurately predicted, demonstrating a potential identifiability issue. However, the average percent deviation from the true rates across a posterior set of 1000 samples remains low, below 40%, for all reactions except reaction 5 which overestimated the rate six-fold, as shown in Table3. Reactions 1, 2, and 6 did particularly well with an average percent deviation of 1.22% ± 0.81, 9.74% ± 0.64, and 4.08% ± 1.28 respectively.

**Figure 16.**
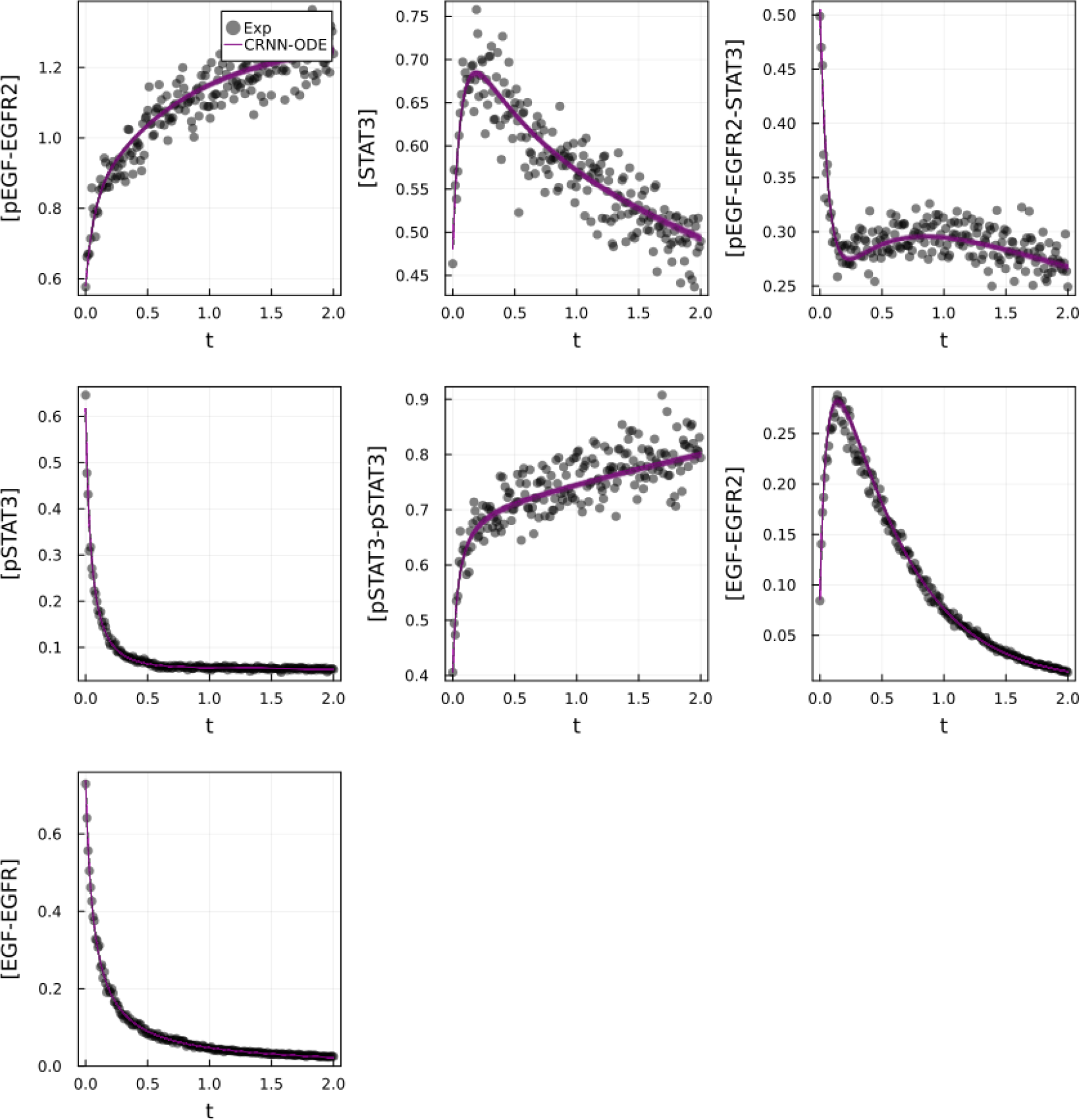
Comparison of Bayesian chemical reaction neural network prediction compared to training data for EGFR-STAT3 pathway described in case 2. Gaussian noise with standard deviation of 5% was added to the training data. 500 posterior sample predictions are superimposed on the data.

**Figure 17.**
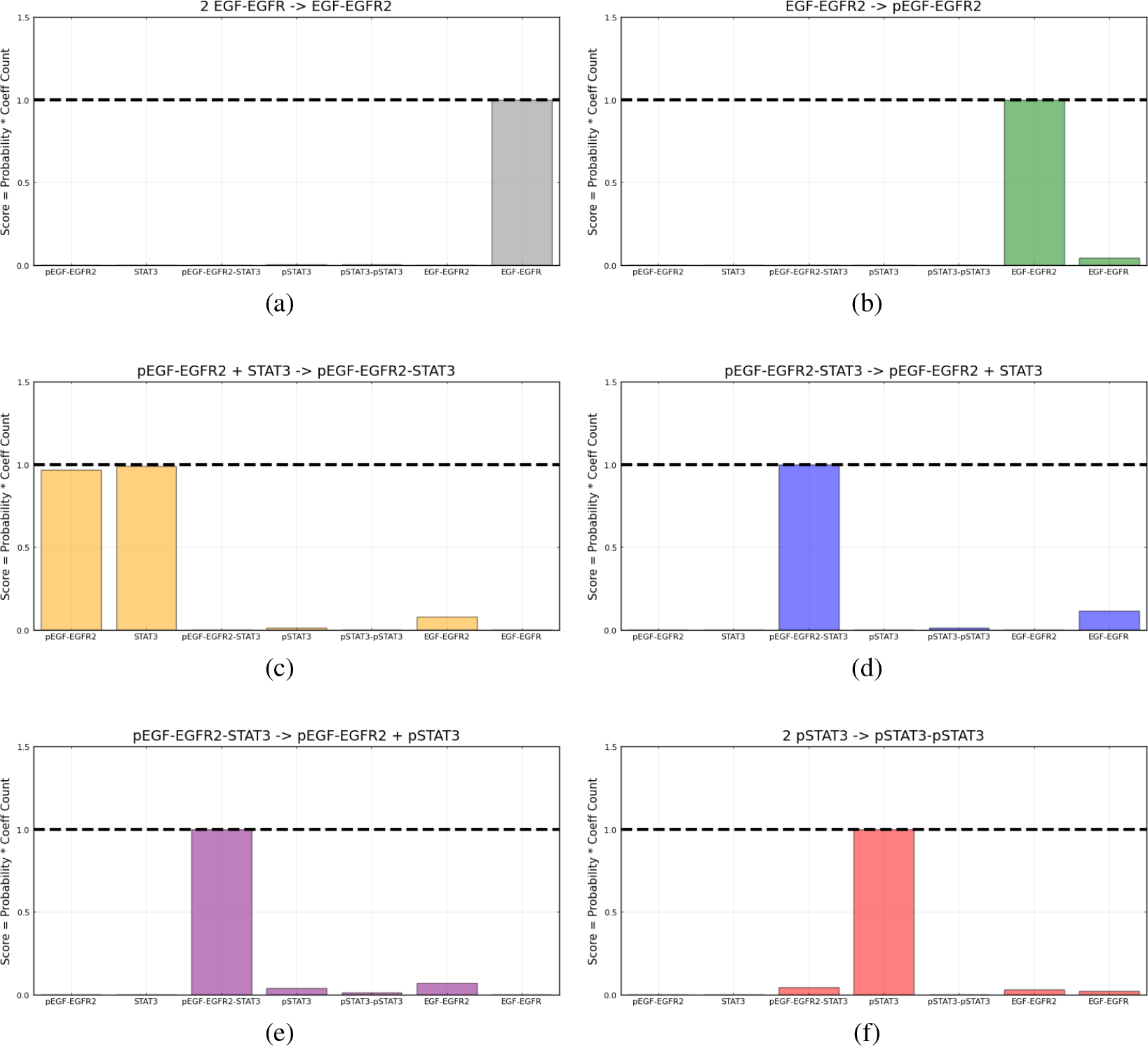
Recovered score metric for each of the six reactions included in case 2. Score metrics are calculated using equation 6 and represent the weighted probability that a species is a reactant of each reaction. Gaussian noise with standard deviation of 5% was added to the training data.

**Figure 18.**
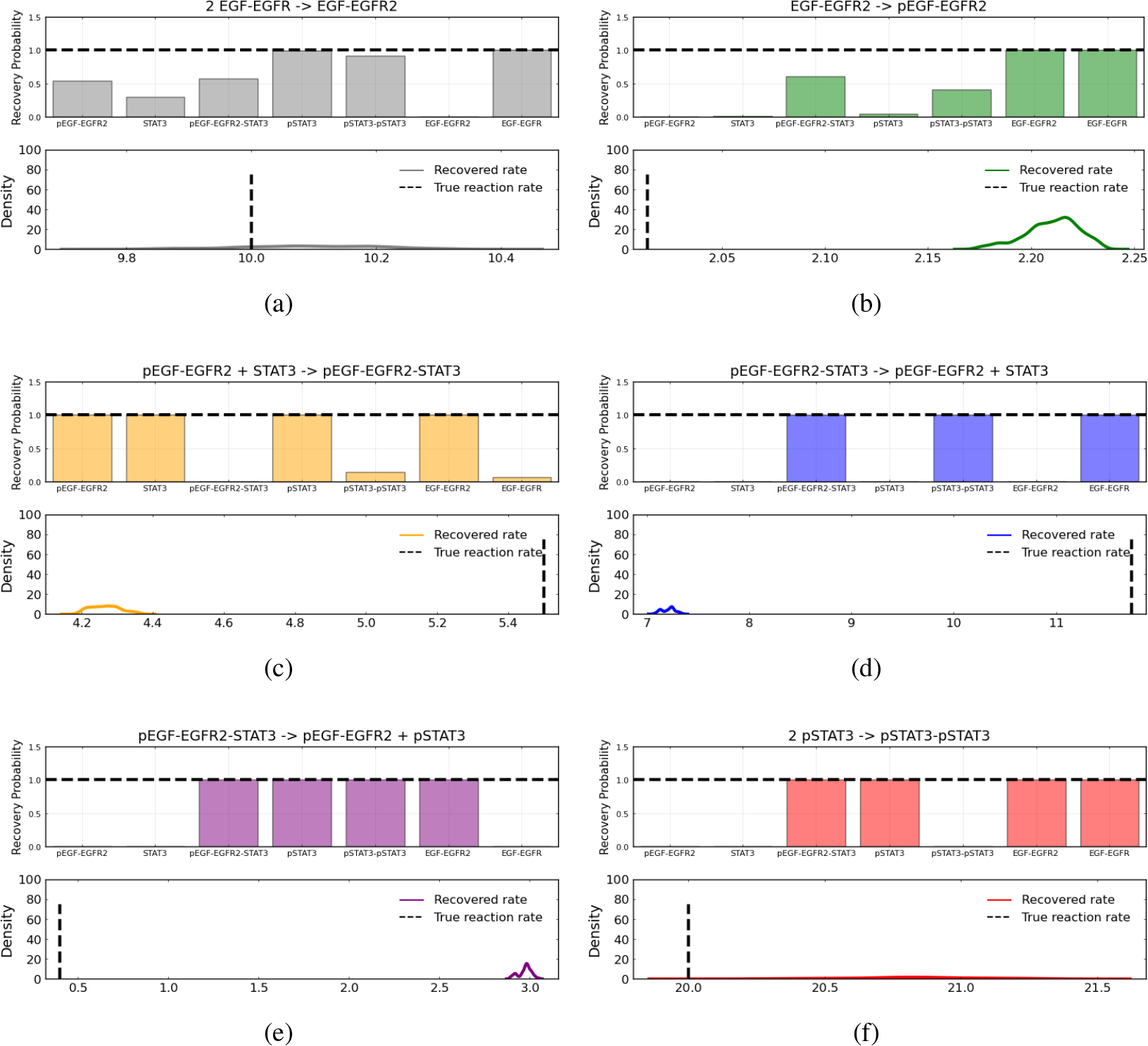
Reactant recovery probability for each of the seven species included in case 2 and posterior distributions of the learned reaction rates for each of the case 2 reactions. Gaussian noise with standard deviation of 5% was added to the training data. A posterior set of 1000 samples was chosen for the estimation.

## 5 Discussion

In this study, we combine the previously published chemical reaction neural network (CRNN) [3] with a preconditioned Stochastic Gradient Langevin Descent (pSGLD) Markov Chain Monte Carlo (MCMC) stepper [21] of neural ODEs, allowing for the efficient discovery of chemical reaction networks from concentration time course data and quantification of the uncertainty in learned parameters. The specialized form of the CRNN is constrained to satisfy the kinetic equations of reactions, which enables the identification of the learned neural network parameters as the stoichiometric coefficients and reaction rates of the reactions while the Bayesian optimizer provides an estimate of the uncertainty of those parameters.

We tested the algorithm with a simple system of four reactions and five species. With no noise added to the training data, the algorithm perfectly captured the reaction network and accurately matched the time course for each species. Additionally, the posterior distributions of the reaction rates were centered around the true reaction rates. With the addition of noise (standard deviation of 5% of the true concentration) to the training data, the reaction networks were also accurately captured, however the posterior distributions of the reaction rates were no longer centered at the true value but still contained within the distribution. Even with the addition of a large amount of noise (standard deviation of 50%), the reactants of each reaction were correctly identified, and posterior distributions still contained the true reaction rates.

We demonstrated that the preconditioned SGLD is necessary to increase training robustness by comparing the results of our Bayesian CRNN trained with the preconditioned SGLD optimizer to a Bayesian CRNN trained with a standard SGLD optimizer and found that not only is the algorithm more efficient and requires less epochs to train but it also more accurately approximates the posterior. The standard SGLD shows high confidence in the incorrect value while the pSGLD in general shows lower confidence but includes the true values.

Additionally, we demonstrated that the Bayesian CRNN in addition to being interpretable, improves upon prediction accuracy when extrapolated beyond the time range used for training by comparing the CRNN to a purely data-driven neural ODE model and an LSTM-based model that contain no prior scientific knowledge about the structure of chemical reactions.

Application of this model to a larger system, a simplified representation of the EGFR-STAT3 pathway containing seven species and six reactions revealed the limitations of this model. While the model was able to accurately predict the time course of all species and learn the correct reactions, as evaluated by correct prediction of reactants and products, the posterior distributions of the reaction rates for four out of six reactions did not include the true rates. This is not completely unexpected since as reaction networks become larger, different combinations of reaction parameters may result in the same concentration dynamics. Future work will be needed to solve this identifiability issue, allowing for the use of this model on larger reaction systems.

The combination of a preconditioned SGLD optimizer with the CRNN allows for efficient training and posterior sampling as well as reliable estimates of parametric uncertainty while accounting for epistemic uncertainty in a way that builds confidence in the learned reaction network and can help determine if more data is needed. This work demonstrates that knowledge-embedded machine learning techniques via SciML approaches may greatly outperform purely deep learning methods in a small-medium data regime that is common in Quantitative Systems Pharmacology (QSP) and demonstrates viable techniques for the automated discovery of QSP models directly from timeseries data.

## References

[1] Christopher Rackauckas, Yingbo Ma, Julius Martensen, Collin Warner, Kirill Zubov, Rohit Supekar, Dominic Skinner, Ali Ramadhan, and Alan Edelman. Universal differential equations for scientific machine learning, 2021.

[2] Raj Dandekar, Chris Rackauckas, and George Barbastathis. A machine learning-aided global diagnostic and comparative tool to assess effect of quarantine control in covid-19 spread. Patterns, 1(9):100145, 2020.

[3] Weiqi Ji and Sili Deng. Autonomous discovery of unknown reaction pathways from data by chemical reaction neural network. The Journal of Physical Chemistry A, 125(4):1082–1092, 2021.

[4] Ricky TQ Chen, Yulia Rubanova, Jesse Bettencourt, and David K Duvenaud. Neural ordinary differential equations. In Advances in neural information processing systems, pages 6571–6583, 2018.

[5] Qiaofeng Li, Huaibo Chen, Benjamin C. Koenig, and Sili Deng. Bayesian chemical reaction neural network for autonomous kinetic uncertainty quantification. Phys. Chem. Chem. Phys., 25:3707–3717, 2023.

[6] Matthew D Hoffman and Andrew Gelman. The no-u-turn sampler: adaptively setting path lengths in hamiltonian monte carlo. J. Mach. Learn. Res., 15(1):1593–1623, 2014.

[7] Max Welling and Yee W Teh. Bayesian learning via stochastic gradient langevin dynamics. In Proceedings of the 28th international conference on machine learning (ICML-11), pages 681–688, 2011.

[8] Tianqi Chen, Emily B. Fox, and Carlos Guestrin. Stochastic gradient hamiltonian monte carlo, 2014.

[9] David J. Lunn, Nicky Best, Andrew Thomas, Jon Wakefield, and David Spiegelhalter. Bayesian analysis of population pk/pd models: General concepts and software. Journal of Pharmacokinetics and Pharmacodynamics, 29(3):271–307, Jun 2002.

[10] Udo von Toussaint. Bayesian inference in physics. Rev. Mod. Phys., 83:943–999, Sep 2011.

[11] Mark Girolami. Bayesian inference for differential equations. Theoretical Computer Science, 408(1):4 –16, 2008. Computational Methods in Systems Biology.

[12] Hanwen Huang, Andreas Handel, and Xiao Song. A bayesian approach to estimate parameters of ordinary differential equation. Computational Statistics, 35(3):1481–1499, Sep 2020.

[13] Laurent Valentin Jospin, Wray L. Buntine, F. Boussaid, Hamid Laga, and M. Bennamoun. Hands-on bayesian neural networks - a tutorial for deep learning users. ArXiv, abs/2007.06823, 2020.

[14] Wesley Maddox, Timur Garipov, Pavel Izmailov, Dmitry P. Vetrov, and Andrew Gordon Wilson. A simple baseline for bayesian uncertainty in deep learning. CoRR, abs/1902.02476, 2019.

[15] Pavel Izmailov, Wesley J. Maddox, Polina Kirichenko, Timur Garipov, Dmitry P. Vetrov, and Andrew Gordon Wilson. Subspace inference for bayesian deep learning. CoRR, abs/1907.07504, 2019.

[16] Raj Dandekar, Karen Chung, Vaibhav Dixit, Mohamed Tarek, Aslan Garcia-Valadez, Krishna Vishal Vemula, and Chris Rackauckas. Bayesian neural ordinary differential equations. arXiv preprint arXiv:2012.07244, 2020.

[17] Christopher Rackauckas, Alan Edelman, Keno Fischer, Mike Innes, Elliot Saba, Viral B Shah, and Will Tebbutt. Generalized physics-informed learning through language-wide differentiable programming. In AAAI Spring Symposium: MLPS, 2020.

[18] Christopher Rackauckas and Qing Nie. Differentialequations. jl–a performant and feature-rich ecosystem for solving differential equations in julia. Journal of Open Research Software, 5(1), 2017.

[19] Hong Ge, Kai Xu, and Zoubin Ghahramani. Turing: a language for flexible probabilistic inference. In International Conference on Artificial Intelligence and Statistics, AISTATS 2018, 9-11 April 2018, Playa Blanca, Lanzarote, Canary Islands, Spain, pages 1682–1690, 2018.

[20] Kai Xu, Hong Ge, Will Tebbutt, Mohamed Tarek, Martin Trapp, and Zoubin Ghahramani. Advancedhmc. jl: A robust, modular and efficient implementation of advanced hmc algorithms. 2019.

[21] Chunyuan Li, Changyou Chen, David Carlson, and Lawrence Carin. Preconditioned stochastic gradient langevin dynamics for deep neural networks. arXiv preprint arXiv:1512.07666, 2015.

[22] Dominic P Searson, Mark J Willis, and Allen Wright. Reverse engineering chemical reaction networks from time series data. arXiv preprint arXiv:1412.6346, 2014.

[23] Mike Innes. Flux: Elegant machine learning with julia. Journal of Open Source Software, 3(25):602, 2018.

[24] Gholamreza Bidkhori, Ali Moeini, and Ali Masoudi-Nejad. Modeling of tumor progression in nsclc and intrinsic resistance to tki in loss of pten expression. PLoS One, 2012.

